# A variational autoencoder trained with priors from canonical pathways increases the interpretability of transcriptome data

**DOI:** 10.1101/2023.05.22.541678

**Authors:** Bin Liu, Bodo Rosenhahn, Thomas Illig, David S. DeLuca

## Abstract

Interpreting transcriptome data is an important yet challenging aspect of bioinformatic analysis. While gene set enrichment analysis is a standard tool for interpreting regulatory changes, we utilize deep learning techniques, specifically autoencoder architectures, to learn latent variables that drive transcriptome signals. We investigate whether simple, variational autoencoder (VAE), and beta-weighted VAE are capable of learning reduced representations of transcriptomes that retain critical biological information. We propose a novel VAE which utilizes priors from biological data to direct the network to learn a representation of the transcriptome that is based on understandable biological concepts.

After training five different autoencoder architectures on 22310 transcriptomes, we benchmarked their performance on organ and disease classification tasks on separate selection of 5577 test samples. Every tested architecture succeeded in reducing the transcriptomes to 50 latent dimensions, which captured enough variation for accurate reconstruction. The simple, fully connected autoencoder, performs best across the benchmarks, but lacks the characteristic of having directly interpretable latent dimensions. The beta-weighted, prior-informed VAE implementation is able to solve the benchmarking tasks, and provide semantically accurate latent features equating to biological pathways.

This study opens a new direction for differential pathway analysis in transcriptomics with increased transparency and interpretability.

**Author summary:** The ability to measure the human transcriptome has been a critical tool to studying health and disease. However, transcriptomes data sets are too large and complex for direct human interpretation. Deep learning techniques such as autoencoders are capable of distilling high-level features from complex data. However, even if deep learning models find patterns, these patterns are not necessarily represented in a way that humans can easily understand. By bringing in the prior knowledge of biological pathways, we have trained the model to “speak the language” of the biologist, and represent complex transcrtomes, in simpler concepts that are already familiar to biologists. We can then apply the tool to compare for example samples from lung cancer cells to healthy cells, and show which biological processes are perturbed.

## Introduction

Transcriptomics is a powerful tool in characterizing cellular activity under various conditions, which allows researchers to discover the underlying associations between transcripts or genes and pathological or environmental factors. Therefore, transcriptomics data are widely applicable in multiple areas of biomedical research, varying from understanding disease mechanisms [1], detecting biomarkers [2, 3], to tissue-specific regulatory gene identification [4]. The broad application of the technology leads to generating a considerable amount of transcriptomic sequencing data and constructing a few specific public platforms hosting the relevant biological data sets, such as ArrayExpress [5] and NCBI GEO [6]. In all these biomedical tasks, human-understandable interpretation of the experimental transcriptomics data serves as the pivotal component to understanding the underlying biology. However, this interpretation remains a challenge in the face of large and complex data sets. Here we explore the potential for machine learning models to learn a simplified representation of the transcriptome in terms of commonly understood biological processes, and thus increase the interpretability.

While gene set enrichment analysis (GSEA) [7] is a standard tool for interpreting regulatory changes to the transcriptome, the method highly relies on a list of well-detected differentially expressed genes (DEGs) between the conditions of interest. Furthermore, most of the state-of-the-art models for differential expression analysis (DEA) are based on the linear assumption across samples, the representatives of which include the models from limma [8, 9], DESeq2 [10], and Seurat [11]. However, variation in measured expression levels, whether of biological or technical origin, may not always behave linearly. Given the potential for synergistic effects between genes, this assumption of linearity might lead to a loss of power. Complex sources of variation are also present in combined public data sets, consisting of a large number of samples from multiple sources, and influenced by non-biological factors such as batches or processing centers. These limitations daunt the exploration of large-scale transcriptomics data for answering fundamental biological questions.

The goal of this study is to utilize deep learning techniques to bring transparency into which biological process patterns are represented in a transcriptome data set, and thus to facilitate the interpretation of experimental results. Deep learning with artificial neural networks has witnessed rapid development in recent years and outperformed the traditional approaches in handling massive and complex data from multiple areas because of its high-level feature extraction at a non-linear space and objective data processing [12–16]. This development also enables more possibilities in understanding the transcriptomics data and has successfully contributed, for example to drug repurposing and development [17, 18], phenotype classification [19] and genomics functional characterization [20] with a more in-depth understanding of transcriptomics data.

We are specifically interested in autoencoders as a class of methods which can reduce the dimensionality and learn the major features in complex data [21]. Autoencoders achieve this by passing the data through a bottleneck layer (a layer with fewer nodes than in input layer) and optimizing the model with the objective of generating an output that is as similar as possible to the input. These methods have been explored in the context of transcriptomes in a few representative publications, including [20, 22–26], demonstrating that interesting biological features are captured in the latent space of the model. However, these studies take different approaches to the challenge of associating latent representations with human-understandable biological concepts. [22] implemented an autoencoder and interpreted that latent space by correlating latent features with phenotypes post hoc. [20] on the other hand sought to constrain the latent space to represent known pathways by restricting the network connectivity, i.e., each latent node represents a pathway, and only genes known to be involved in that pathway are connected to the node upstream. In the study of [22], the network is free to learn any representation from that data, but the burden of interpretation is left to post hoc analysis. In the case of [20], the network is restricted directly in its architecture based on gene set definitions.

Here, we see an opportunity to implement a solution that finds a middle path in which the network is encouraged to learn a latent representation based on known biological concepts, but still has the freedom to learn relationships among genes from the data. Specifically, we propose an autoencoder variant using a novel technique of introducing pathway-informed priors. The basis for this approach is the Bayesian framework implemented in variational autoencoders (VAE) [27]. The VAE framework inherently provides the opportunity to involve priors in the training process to learn latent representations. With the introduction of biologically meaningful priors, our approach here aims to integrate prior knowledge from the domain (here, we use Hallmark pathways defined by MSigDB as an example) and still retain the flexibility of the data-driven deep learning approach. This approach is most comparable to those presented by Zhao [25] and Lotfollahi [26], with the commonality that the goal is to produce latent features that correspond to known biological concepts. However, compared to our approach of incorporating canonical pathway knowledge as priors, both Zhao [25] and Lotfollahi [26] provide pathway definitions to constrain the decoder architecture. The prior-based approach provides an additional opportunity to calibrate the strength of the effect of prior following the established beta-VAE approach ([28]).

Specifically, We make use of a hyperparameter, beta, in a similar way described by [28], which can add weight to the influence of the priors on the training solution. Thus, using the beta, we can control the extent to which the model conforms to pathway concepts previously defined by MSigDB versus being free to define latent variables in any way that best encodes the transcriptome in a reduced representation. The fine-tuning of this hyperparameter enables control over the tension between direct biological interpretability and the ability to deviate from canon or find new patterns.

In this study, we implement and compare several standard autoencoder implementations: (i) fully connected autoencoder (simpleAE) [29], (ii) variational autoencoder (simpleVAE), (iii) beta-VAE (beta-simpleVAE), as well as (iv) novel derivatives using prior biological data: priorVAE, and (v) beta-priorVAE. In order to benchmark the performance of the series of models proposed here, we perform tissue and disease classification (e.g., adenocarcinoma, small cell lung cancer (SCLC)) based on the latent variables discovered in the models. This study explores the feasibility of the prior-based VAE approach in increasing the transparency and interpretability of transcriptomes, as well as how the hyperparameter beta controls the balance of using prior information versus learning novel patterns directly from the data.

## Materials and methods

### Data sets and preprocessing

The data set employed for the training of the model and subsequent analyses was downloaded from ArrayExpress [5], with Accession ID E-MTAB-3732 [30]. This data set comprises 27,887 Affymetrix HG-U133Plus2 arrays, sourced publicly. All samples underwent quality control filtering and were annotated for disease status and cell line information. The data was normalized using fRMA [31–34] within the R Bioconductor platform by the data set’s author. The data set contains samples from healthy individuals, those with diseases (including cancer), and cell lines. The original data set has been divided into a training and a test set at an 8:2 ratio, with stratification based on the source organs of the samples. The curated gene signature data set, used in the previous generation, is the Hallmark gene set, sourced from Human MSigDB Collections [35].

The transcriptome data were prepared as input into the autoencoders in three variations. In the first input variation, the data were fed into the model on the transcriptome level without further processing. For the second input variation, the transcripts were collapsed into the gene level using the platform annotation offered by Gene Expression Omnibus (GEO) [6]. The normalized expression levels of the transcripts were transformed into the original level by applying a power function. The original expressions were averaged, and a log2 transformation was processed on the mean values.

For the third input variation, the goal was to decrease the number of trainable parameters in the models further. A community detection algorithm was applied to the gene expression level. An unsupervised nearest neighbors learning was applied to the absolute level of correlation between each pair of genes using the NearestNeighbors function from the sklearn Python package [36]. A graph of k-Neighbors was computed to make a graph of gene relationships based on the expression levels. We then use the Leiden algorithm [37] to detect the communities in this graph. A total of 2032 communities were defined using a resolution value of 0.02. A single gene was chosen as a representative of each community for use as the final input. Community representatives were based on the criteria of having the highest sum of correlation with all the other genes in the same community.

### Model architectures

The experiment systematically trained and compared five architectures of autoencoders, including one fully-connected autoencoder without prior information (simpleAE) and four variational autoencoders (VAE) architectures. Besides the variational autoencoder with unit Gaussian prior (simpleVAE) and the beta-constrained variational autoencoder with unit Gaussian prior (beta-simpleVAE), we also presented a novel technique of introducing pathway-informed priors (priorVAE) and tested the influence of the hyperparameter beta over this biological relevant prior VAE (beta-priorVAE).

We construct three fully-connected linear layers in the encoder of all the autoencoder architectures (number of trainable parameters equal to the dimension of features, 1000 and 100, respectively). The leaky rectified linear unit (Leaky ReLU) serves as the activation function between encoder layers.

The bottleneck layer of the simpleAE is a dense layer with 50 dimensions. For the VAEs, the bottleneck is implemented as two fully-connected, 50-dimension layers: one for learning *σ*s and one for learning *µ*s, which are then sampled using the reparametrization trick before going on to the decoder. The decoders in all models mirror the three dense layers of the encoders. The Softplus activation function is added to the last layer of the decoder for reconstruction to output values on the same scale as gene expression input values.

The loss function for the simpleAE is the mean squared error (MSE) between the decoder output and the original input values. For VAE, the loss function has two terms: (i) the reconstruction loss, also MSE, and (ii) the KL divergence between the latent distributions and unit Gaussian prior distributions. For the prior- and beta-priorVAE implementations, the loss functions are described in detail below.

### The preparation of pathway-informed priors

Pathway-informed prior distributions were generated for each sample in the form of (*σ*^2^) and (*µ*) parameters using a bootstrapping procedure. We defined *µ* as the average gene expression level of genes within each pathway definition, and *σ* as the variance across bootstrapping iterations. Pathway definitions were taken from MSigDB Hallmarks [35].

### Loss function for biological priors

Generally, variational autoencoders take the form found in Fig 1 and have been previously described in detail [27]. In short, it has been established that neural network training can perform variational inference when the loss function takes the form:

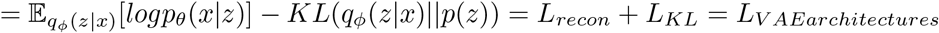

**Fig 1.**
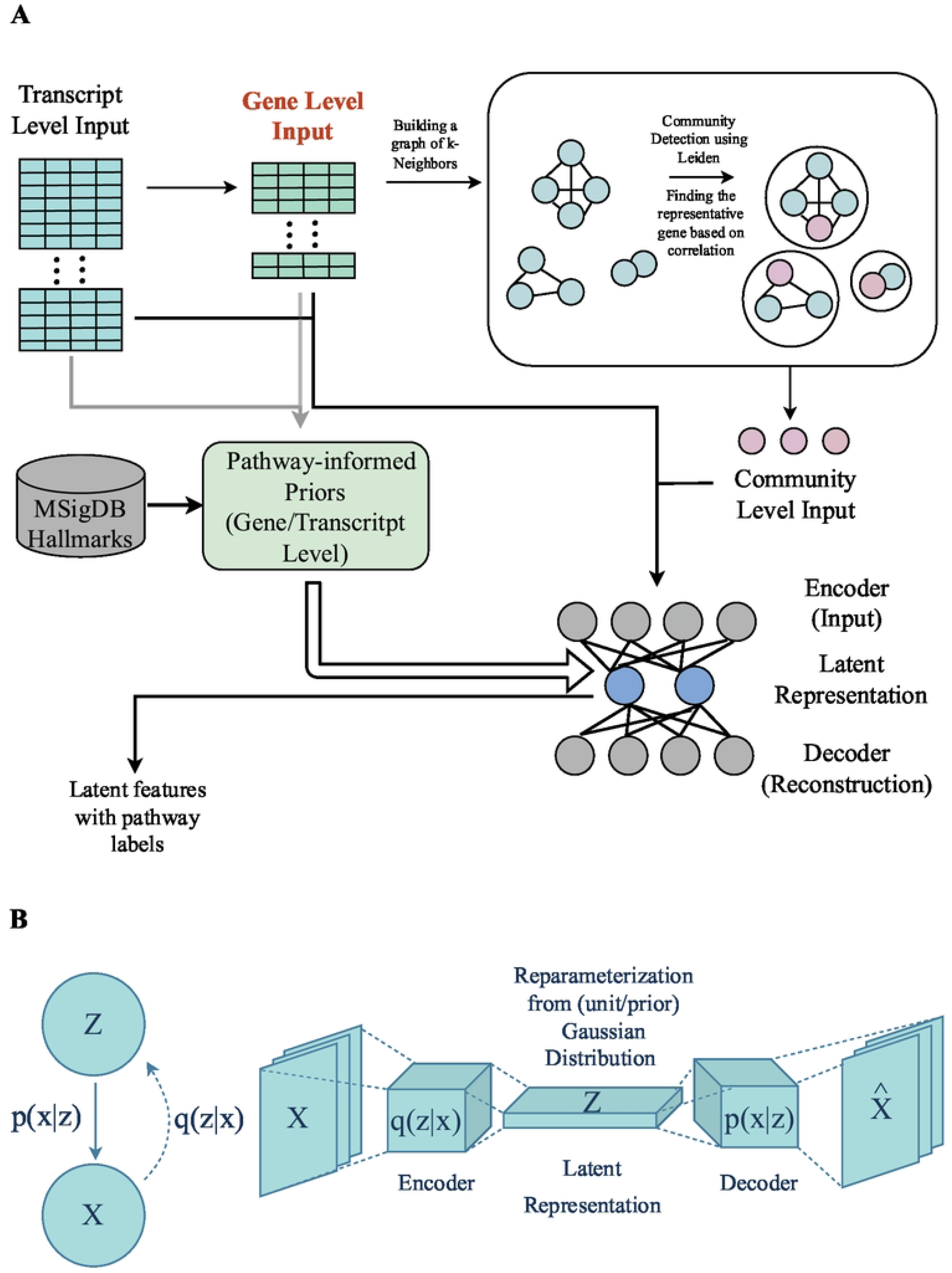
A conceptual illustration of variations autoencoder architectures. A: An overview of the system architecture with different input levels. A pathway-derived prior is generated on the transcript or gene level, as described in the following sections. The training and post hoc are repeated with input on the transcript, gene, and community levels. B: As a latent variable framework, the model assumes that latent variables (*Z*) are determinants of the measured data (*X*). To learn *Z, p*(*Z*| *X*) is approximated by *q*(*Z*|*X*), and modeled as the encoder portion of the network. The latent values represent probability distributions, which are implemented in the bottle-neck layer as values for mu (*µ*) and sigma (*σ*). Finally, the decoder is conceptually equivalent to *p*(*X*|*Z*).

Here, *x* ∈ ℝ is a vector of expression values for a sample in the set of all samples, *X*. The generative model *p*_*θ*_(*x*|*z*) is learned, where *z* (*z* ∈ ℝ) are latent variables with a prior *p*(*z*) such that *z* can generate the observed data *x*.

The reconstruction loss term *L*_*recon*_ can be measured by the mean squared error (MSE) between the input and the reconstructed output. In most VAE implementations, a Gaussian distribution is used for *p*(*z*) to make a tractable KL calculation. The assumption of unit Gaussian, 𝒩(0, *I*), as priors leads to the simplified expression:

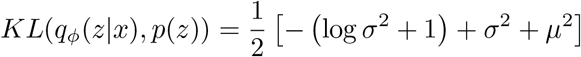

In order to implement pathway-informed priors with any parameter values, the KL divergence had to be implemented more generically for any two Gaussian distributions to measure the distance between *q*_*ϕ*_(*z*|*x*) and *p*(*z*), where 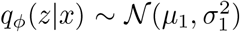 and 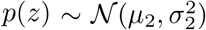

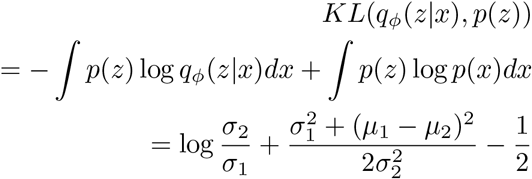

The models beta-simpleVAE and beta-priorVAE involve the addition of hyperparameter *β* (*β >* 1) to put more weight on the *L*_*KL*_ term, as described in [28]:

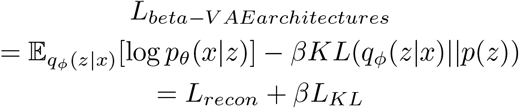

## Results

### Input layer

The Affymetrix HG-U133Plus2 chip captures expression at the transcript level, providing an input space of roughly 50,000 features. Concerned that this would result in a parameter space that was too large for the available data, we experimented with collapsing the inputs to the gene level and even further to ‘representative genes’ using a network-based clustering technique. A comparison of the choice of inputs can be found in S1 Fig. We concluded that gene-level input was a sufficient reduction in the parameter space based on the results. The results reported below are based on this gene-level input.

### Learning latent representations with several autoencoder architectures

Five autoencoder variations were trained on the full set of transcriptomes provided by the ArrayExpression dataset, E-MTAB-3732, containing 27887 samples. As shown in Table 1, common to each architecture is a 50-dimensional latent space. For the models, simpleAE, simpleVAE, and beta-simpleVAE the 50 latent dimensions were learned strictly from the data, without attributing any prior biological concepts, and are simply enumerated 1 through 50. For the priorVAE and beta-priorVAE, the 50 latent nodes are associated with the 50 pathways found in the MSigDB Hallmarks gene sets, and accordingly, each latent node can be labeled with the gene set name. To evaluate the training results, we use three types of benchmarking strategies, as seen in Table 2. (i) an evaluation of reconstruction performance, (ii) the Kullback–Leibler (KL) divergence between the latent distributions and the prior distributions, and (iii) the combined total loss.

**Table 1.**
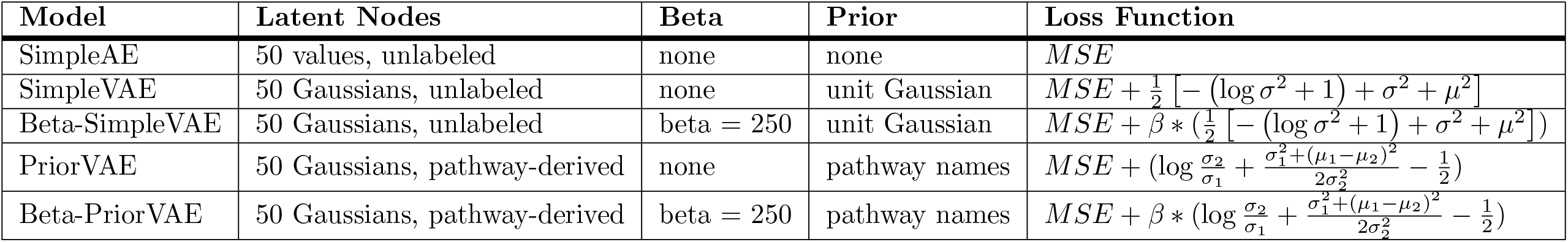
An overview of the structures of the five architectures.

**Table 2.** Benchmark results for the five architectures on the test set.

| Model | Recon. Loss | KL(w.β) | KL(n.β) | Total Loss (w.β) |
| --- | --- | --- | --- | --- |
| SimpleAE | 2674.6 | NA | NA | NA |
| SimpleVAE | 2780.8 | 184.5 | 184.5 | 2965.3 |
| Beta-SimpleVAE | 5899.7 | 7.4 | 1846.8 | 7746.5 |
| PriorVAE | 2758.5 | 160.3 | 160.3 | 2918.8 |
| Beta-PriorVAE | 5740.2 | 8.1 | 2012.9 | 7753.1 |

Loss function values broken down into reconstruction loss, KL divergences with $\beta$ (w.β), without $\beta$ (n.β)

Taking these metrics together, it is clear that the introduction of hyperparameter *β* results in a lower KL divergence at the cost of a slightly worse reconstruction loss. The introduction of priors resulted in little change to the loss terms compared to their non-prior counterparts.

To further explore reconstruction performance, we computed correlation coefficients between input transcriptomes vs. the output transcriptomes. A complete pairwise correlation analysis across samples shows how similar input and output transcriptomes are to each other in the context of the natural variability across samples (Fig 2). Each model was able to reproduce reasonable output transcriptomes, which correlated with R between ***0*.*97 and 0*.*8***. In most cases, the closest pairwise correlations were between the input and output models, although this was not the case for many samples in the beta-simpleVAE, and a single sample in the beta-priorVAE. Together with the performance shown in Fig 2, it is clear that increasing the beta hyperparameter and emphasizing the influence of the priors comes at a cost to reconstruction performance.

**Fig 2.**
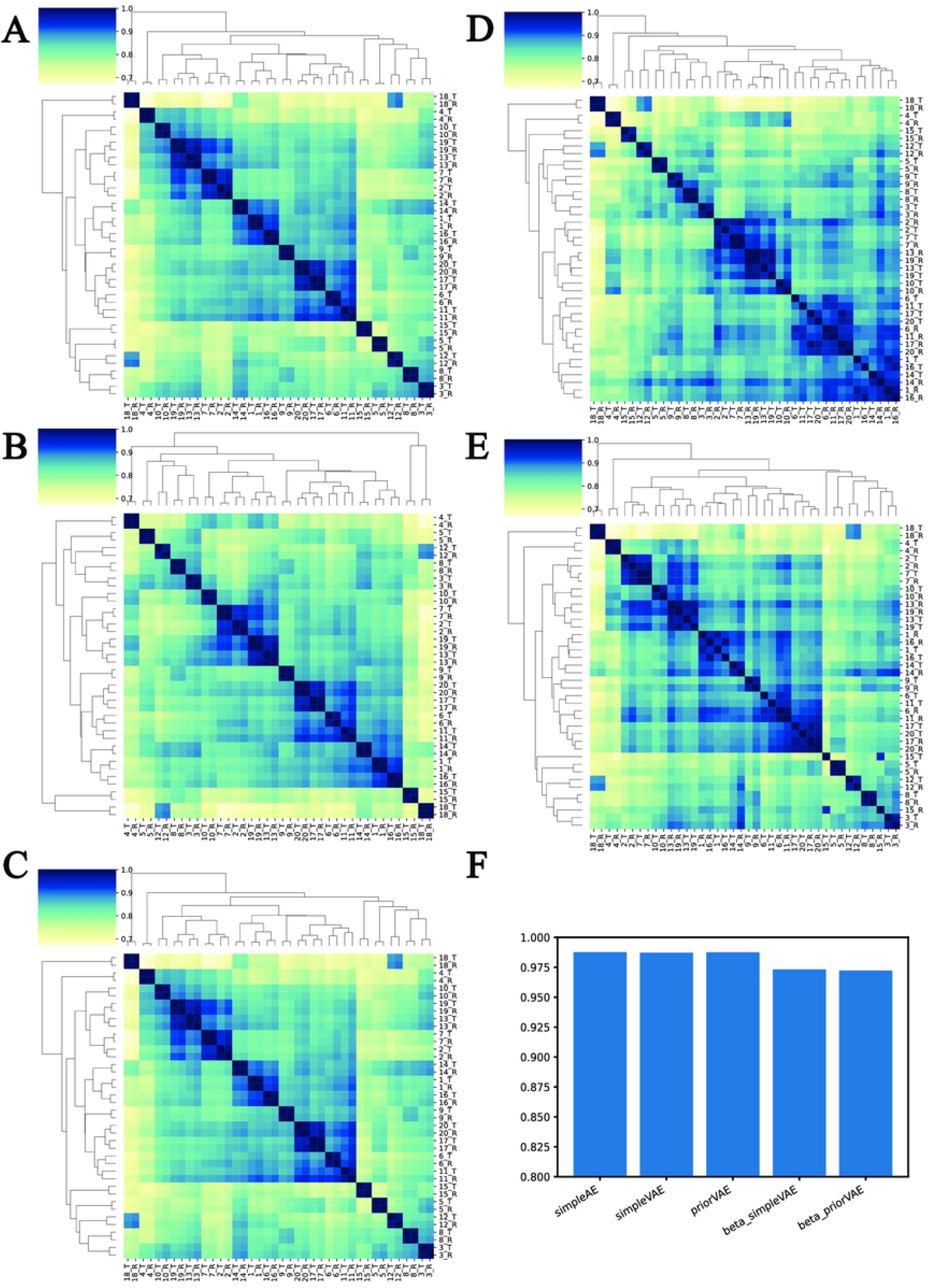
Reconstruction performances using correlation coefficients between input and output transcriptomes. A-E: The clustered pair-wise correlation heatmaps of the selected input and their reconstructed output for A: simpleAE, B: simpleVAE, C: priorVAE, D: beta-simpleVAE, E: and beta-priorVAE. Selected input samples and their corresponding reconstruction output are enumerated as 1-20. T represents the input train sample and R represents the reconstructed output. F: The average correlation between the input and its corresponding reconstruction output.

Ideally, the latent dimension should contain enough information to capture the essential biological features. To systematically evaluate how well the models can perform in this regard, we have used the latent features as input into classification models for three validation tasks: (i) distinguishing tissue types (ii) distinguishing healthy vs. adenocarcinoma, and (iii) distinguishing adenocarcinoma from small cell lung cancer (SCLC). The classification performance after five-fold cross-validation is reported in Fig 3, 4, and 5.

**Fig 3.**
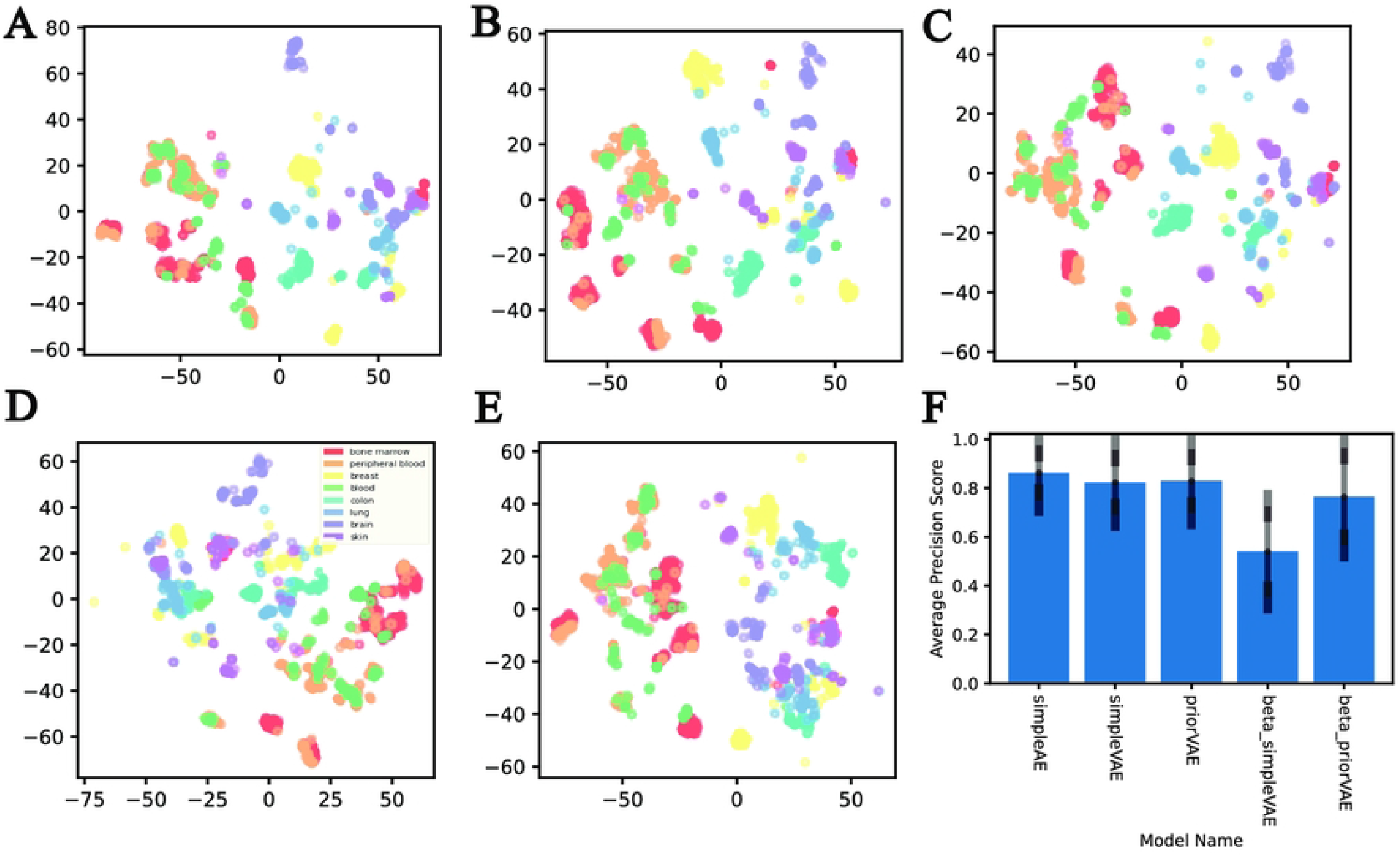
The results and performance of the AE models on the tissue classification. A-E: The t-distributed stochastic neighbor embedding (t-sne) representations of the latent values (*µ*) representation of samples from different tissue origins for A: simpleAE, B: simpleVAE, C: priorVAE, D: beta-simpleVAE, and E: beta-priorVAE. F: The average precision score of the multivariate logistic regression models sourcing from the latent representation of the models across the eight selected tissue types.

**Fig 4.**
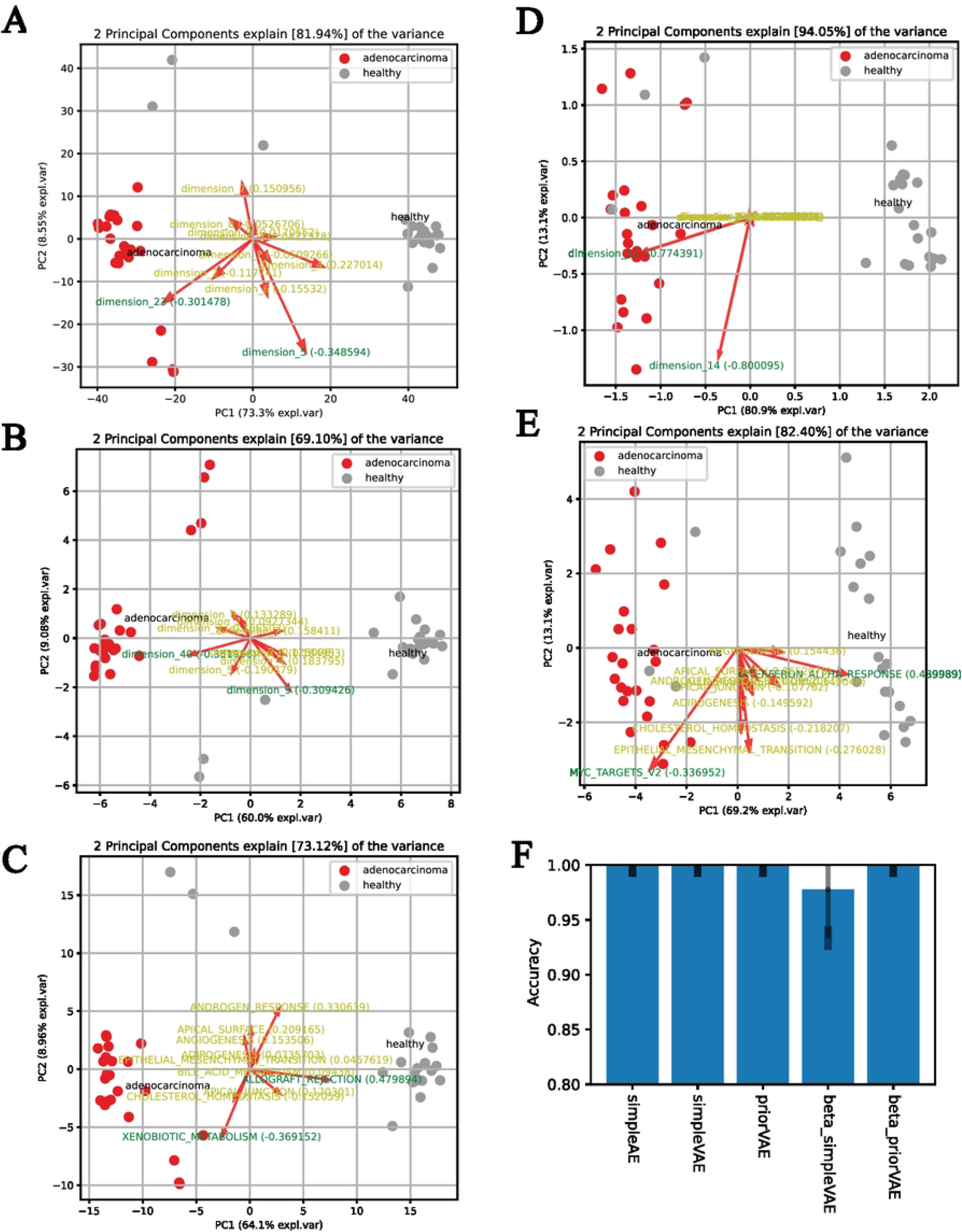
The results and performance of the AE models on the classification of healthy vs. adenocarcinoma samples. A-E: The biplots for the top two principle components (PCs) distinguishing the adenocarcinoma samples and the healthy lung samples for A: simpleAE, B: simpleVAE, C: priorVAE, D: beta-simpleVAE, and E: beta-priorVAE. F: The average precision score of the multivariate logistic regression models sourcing from the latent representation of the models on the task of healthy vs. adenocarcinoma classification.

**Fig 5.**
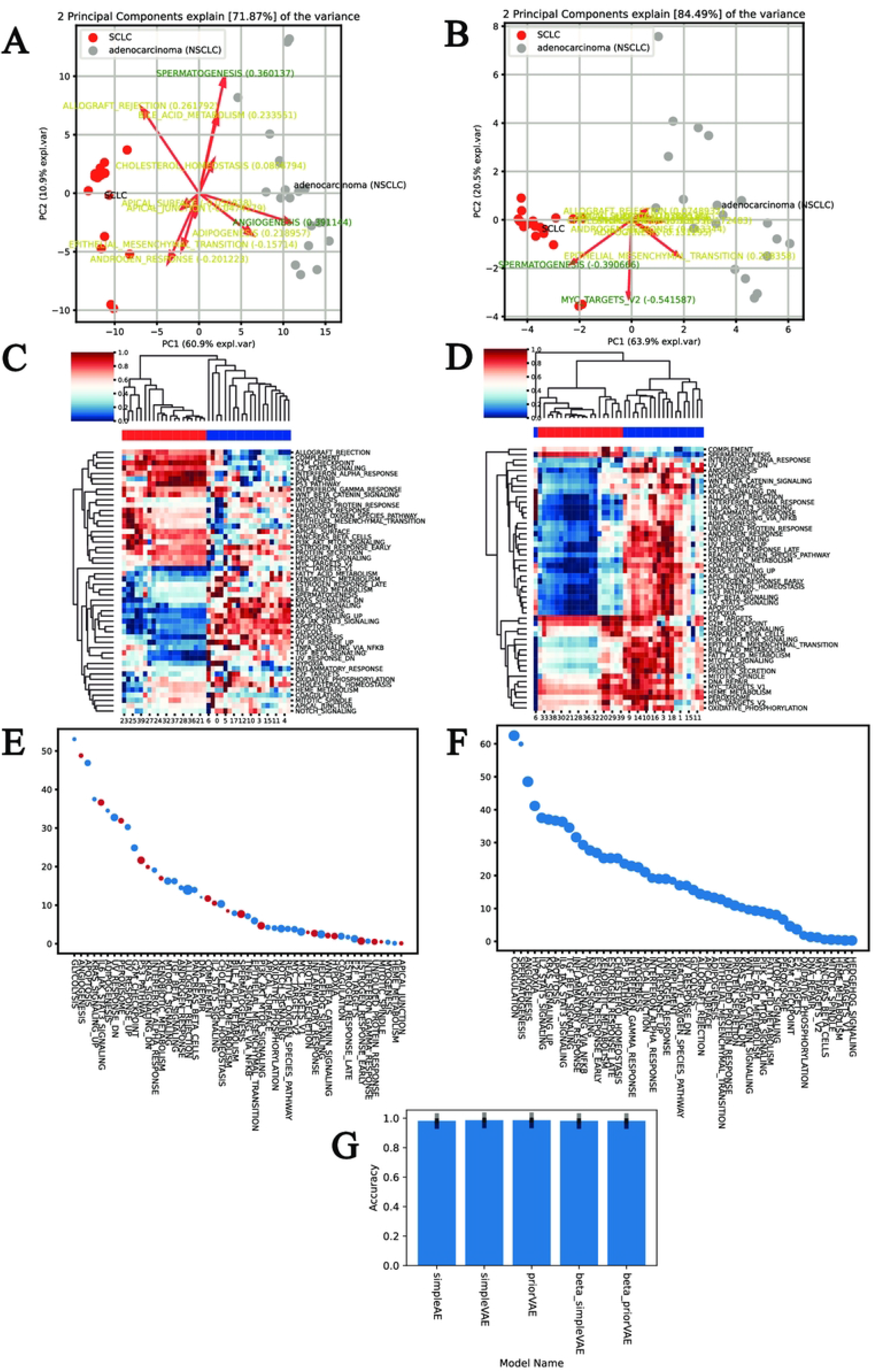
The results of the prior variants (priorVAE and beta-priorVAE) on the classification of lung cancer subtypes (adenocarcinoma vs. small cell lung cancer). A and B: Biplots and principle components analysis (PCA) results of priorVAE and beta-priorVAE. C and D: Heatmaps of the latent results based on selected disease samples for priorVAE and beta-priorVAE. Data is scaled across the fifty latent dimensions. E and F: Two-sided t-test results of the pathway dimensions with the two disease statuses. The significance levels (the p-values) are transformed in the -log10 form and shown on the y-axis. The size of the points represents the correlation between the prior and the latent *µ*, with a larger size indicating a higher level of correlation. The dimensions with a negative correlation are colored in red. G: The average precision score of the multivariate logistic regression models sourcing from the latent representation of the models on the task of adenocarcinoma vs. SCLC classification, showing that each model type is well suited for this classification task.

For the benchmarking task of classifying tissue types, eight target organs were selected based on available samples. In this classification task, the simpleAE, simpleVAE and priorVAE perform with an average precision above 80%. Beta-weighting leads to worse classification performance in both beta-simpleVAE and beta-priorVAE. However, beta-priorVAE performs much better in this validation task than the former (roughly 0.76 vs. less than 0.6).

For the task of distinguishing adenocarcinoma from healthy controls, all five architectures perform comparably well, with precision between 98.6 and 100 percent. Four architectures, (all but beta-simpleVAE) can still distinguish the adenocarcinoma cell line, a subpopulation of the non-small cell lung cancer (NSCLC), from the small cell lung cancer (SCLC) cell lines with 100 percent of average precision. The performance of beta-simpleVAE latent results as a classifier slightly decreased from 98.6% to 97.5%, and the standard deviation (sd) across the five-fold validation increased from 0.016 to 0.049.

### Influence of priors on learned latent representation

The goal of bringing pathway-derived priors into the model is to produce latent representations that both accurately represent the full transcriptome and also directly correspond to recognizable biological concepts. However, these goals are at odds, as we reported in (Fig 6 and S2 Fig). In the case of the priorVAE, the latent variable *allograft rejection* does retain a high correlation with the prior scores (R = 0.82). However, all the other 49 pathways are with a correlation weaker than 0.5, and only six pathways are to some extent correlated with the prior, even with a far less stringent threshold of 0.4. 21 out of 50 pathways correlate negatively with the prior scores. In contrast, the latent variables of the beta-priorVAE retain their connection to their pathways, with 48 of 50 pathways correlating higher than 0.6. Fig 6C indicates that the beta-priorVAE provides latent values with a high level of semantic meaning, providing a direct means for interpreting complex transcriptome data sets.

**Fig 6.**
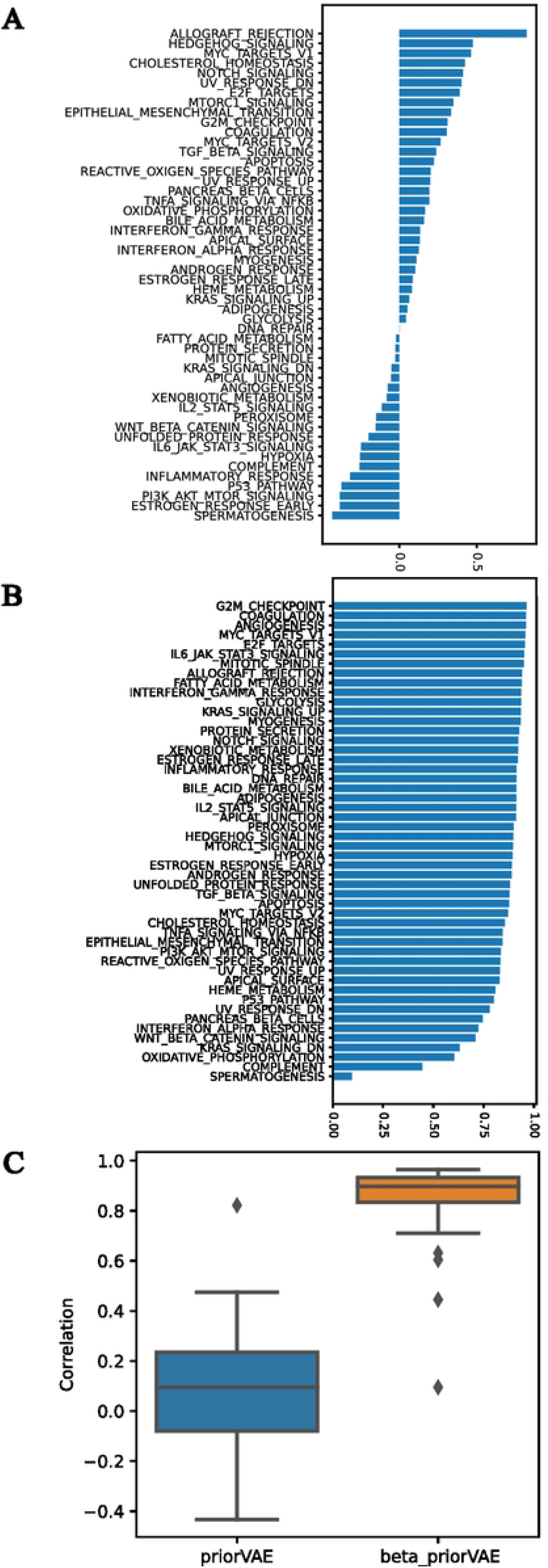
**The semantic meaningfulness of the latent variables in the prior-based models, shown as the correlation between the biological priors and the latent** *µ* **of prior-based models on the test set**. The correlation of each dimension is shown in (A) for priorVAE and (B) for beta-priorVAE. Subplot (C) summarizes these correlations to directly compare the semantic interpretability of the two models.

The beta-priorVAE provides a simplified representation of the transcriptome in terms of 50 features corresponding to 50 pathways. A comparative analysis across samples and conditions is now possible directly on the bases of these features. For the priorVAE model, the biplots of adenocarcinoma and healthy samples indicate that *allograft rejection* and *xenobiotic metabolism* are major features associated with disease (Fig 4). However, for beta-priorVAE *interferon alpha response*, and *MYC targets v2* are the main distinguishing features. In a direct comparison of adenocarcinoma and SCLC, the biplots show that *angiogenesis* and *spermatogenesis* are the two pathways with the highest load to the first principle component (PC) in priorVAE, and *MYC targets v2* and *spermatogenesis* in the case of beta-priorVAE.

A more direct comparison can be made by performing a two-tailed t-test on the values from the two lung cancer phenotype conditions. The results indicate that the top three most differentially expressed dimensions in priorVAE are: *glycolysis, angiogenesis*, and *apoptosis*; for beta-priorVAE, the top three are *coagulation, spermatogenesis*, and *angiogenesis*.

These differences between priorVAE and beta-priorVAE are explained to a large extent by the fact that the beta-priorVAE produces latent variables which track more closely with the pathway-based priors (Fig 6). A direct interpretation of the involvement of a given pathway is only possible when that latent feature is highly correlated with the prior distributions. In the case of adeno vs. small cell, the most distinguishing features, *glycolysis* and *angiogenesis* have only weak correlations with their priors: R = 0.043 and -0.075, respectively. In contrast, for the beta-priorVAE, the most statistically significant feature distinguishing adeno from the small cells is *coagulation*, which is a feature that is also highly correlated with its prior (R=0.96). Therefore, it is only for the beta-priorVAE model that the labels on the latent features retain the meaning of the original pathways. Even for this model, however, it should be noted that the second most statistically significant differentiator is reported as *spermatogenesis*, which is in fact the latent value that has the lowest correlation out of all latent variables for this model (R=0.095). Therefore, it must be concluded that the model detected a feature that is a critical source of variation distinguishing adeno from small cell samples but that this feature is not represented in the set of 50 MSigDB Hallmark pathways.

The comparison between the priorVAE and beta-priorVAE show clearly a trade-off between capturing the biological variability in the models’ latent space, but meanwhile, adhering to prior biological concepts found in the set of Hallmark pathways. To further investigate the effect of hyperparameter beta on performance, we ran the benchmarks across a range of values for beta (S3 Fig) The classification performance seems to decrease consistently with an increasing beta, although for beta values up to 100, the trend is close to flat. This implies that we can find a beta value that balances the need to capture biological features and, at the same time, adhere to the pathway labels provided via the priors.

## DISCUSSION

The results of these experiments demonstrate that autoencoders are capable of generating a simplified representation of a transcriptome that still retains the key biological information necessary to differentiate different cells under different conditions. Furthermore, it is also possible to constrain the training process in a way that forces the network to find a latent representation corresponding to human-understandable biological concepts. Here we have achieved this by taking advantage of the VAE framework, which allows for integrating prior knowledge. There is a trade-off between efficiently representing the complexity of a transcriptome, and adhering to a panel of chosen biological concepts, in our case, defined by 50 Hallmark pathways.

The primary goal of utilizing pathway-based priors in the priorVAE and beta-priorVAE models was to generate a latent space that would be immediately interpretable to a biologist because the model will describe a transcriptome in terms of features that are familiar to a biologist. However, we have observed that latent features do not always retain the identity of their associated priors. In the case of the priorVAE (i.e. beta = 1), the model has substantial freedom to deviate from the pathway priors, and in fact, does so for many features. However, by boosting the requirement to adhere to the pathway concepts with the beta hyperparameter, the immediate biological interpretation of the latent space is achievable (Fig 6C). A notable exception is the case of *spermatogenesis*, which was the feature that had the lowest correlation with the prior. This indicates that the model does require some freedom to discover major sources of variation beyond those pre-defined by the 50 MSigDB Hallmarks. This could indicate a limitation inherent to the 50 Hallmarks, or it could be a limitation of the idea of relying only on known pathways as a source of variation across transcriptomes. There could also be technical sources of variation that need to be accounted for that would not be represented in a pathway database. The presence of unanticipated sources of variation indicates that an interesting future direction for this research would be to include a few “wild-card” nodes in the latent space, with unit-Gaussian priors that are not driven by pathway data. This modification would allow the modeling to account for “unexpected” sources of variation while at the same time utilizing prior pathway information when possible. The burden of interpreting wild-card nodes would be placed on additional *post hoc* analysis.

The beta-priorVAE with the current setting shows an overall satisfying correlation between the latent variables and prior scores, which enables an interpretation of the biological pathways involved in the chosen vignette. Based on the t-test results (Fig 5), *coagulation* is the most significant differentiator between the two lung cancer phenotypes (adenocarcinoma as the representative NSCLC and the SCLC). While coagulation function is associated with the prognosis in NSCLC patients as described in [39, 40], both [41] and [42] reported an absence of similar correlation between coagulation and the SCLC prognosis. The literature is, therefore, consistent with the latent representation in that the concept of coagulation is a differentiating feature between the two diseases. Thus, the evaluation supports the feasibility of such architecture in making transcriptomes more intuitively transparent and interpretable.

Although our primary motivation for including priors in our VAE was to make the latent space directly interpretable, the traditional motivation for including priors was to increase model accuracy. For most of our benchmarks, model accuracy generally decreased when we increased the emphasis on priors via the beta parameter. This implies that the reconstruction portion of the loss function is the main driver of performance in these benchmarks in comparison to the KL divergence term. However, an interesting point of comparison in our experiments is between the beta-simpleVAE and beta-priorVAE, which have the same emphasis on priors (i.e., the same betas), but in the latter model, prior biological knowledge is incorporated. Table 2 and S3 Fig show an increase in performance when using a biological prior vs. a unit Gaussian prior, indicating that in this local comparison, prior biological information can be beneficial.

We have provided evidence that the key sources of biological variation are captured in the latent space. At the primary level, the successful reconstruction as demonstrated by the reconstruction loss, as well as high correlation coefficients between inputs and outputs, indicate that the latent representations are reliable. This is further supported by the fact that cancer types and tissue types can be distinguished using only the latent features. However, the performance for classifying tissues is surprisingly mediocre. The question is whether this is a limitation of the information found in the latent representation or an inadequate classification procedure. The latter scenario is supported by the high reconstruction accuracy and the fact that the multivariate logistic regression maybe be inadequate without proper feature selection.

## Conclusion

The training and post hoc experiments demonstrate that (i) autoencoder models can find simplified representations of transcriptomes that still retain biological information, using pathway-derived priors, we can encourage the models to find latent representations that still adhere to concepts that are familiar to biologists, and (iii) latent features can provide a direct means of comparison among samples and conditions that can provide an immediate biological interpretation. This area of research should be explored further, with attention to alternate pathway definitions to define the priors and thus the latent space, additional model architectures, and integration into bioinformatic workflows.

## Supporting information

**S1 Fig. The performance of reconstruction correlation and the biplots for healthy vs. adenocarcinoma classification on the level of community-, gene- and transcript-level input**.

**S2 Fig. The scatter plot of the priors (x-axis) and the latent** *µ* **(y-axis) of all test samples for A: priorVAE and B: beta-priorVAE**.

**S3 Fig. The average precision score of the beta-simpleVAE and the beta-priorVAE models on the tissue classification with different values for hyperparameter beta**.

## Data availability

The source code for this paper is available on GitHub with the following link: https://github.com/BinLiu9205/deepRNA_autoencoder.git and on figshare with the following link: https://figshare.com/articles/software/deepRNA_autoencoder/22227217

## References

1. Kotula-Balak, M., Duliban, M., Gurgul, A., Krakowska, I., Grzmil, P., Bilinska, B., and Wolski, J. K. (2021) Transcriptome analysis of human Leydig cell tumours reveals potential mechanisms underlying its development. Andrologia, 53(11), e14222.

2. Kim, S. H., Kim, J. H., Lee, S. J., Jung, M. S., Jeong, D. H., and Lee, K. H. (2022) Minimally invasive skin sampling and transcriptome analysis using microneedles for skin type biomarker research. Skin Research and Technology, 28(2), 322–335.

3. Dubois, J., Rueger, J., Haubold, B., Far, R. K.-K., and Sczakiel, G. (2021) Transcriptome analyses of urine RNA reveal tumor markers for human bladder cancer: Validated amplicons for RT-qPCR-based detection. Oncotarget, 12(10), 1011.

4. Consortium, G. (2020) The GTEx Consortium atlas of genetic regulatory effects across human tissues. Science, 369(6509), 1318–1330.

5. Parkinson, H., Kapushesky, M., Shojatalab, M., Abeygunawardena, N., Coulson, R., Farne, A., Holloway, E., Kolesnykov, N., Lilja, P., Lukk, M., et al. (2007) ArrayExpress—a public database of microarray experiments and gene expression profiles. Nucleic acids research, 35(uppl 1), D747–D750.

6. Barrett, T., Wilhite, S. E., Ledoux, P., Evangelista, C., Kim, I. F., Tomashevsky, M., Marshall, K. A., Phillippy, K. H., Sherman, P. M., Holko, M., Yefanov, A., Lee, H., Zhang, N., Robertson, C. L., Serova, N., Davis, S., and Soboleva, A. (11, 2012) NCBI GEO: archive for functional genomics data sets—update. Nucleic Acids Research, 41(D1), D991–D995.

7. Subramanian, A., Tamayo, P., Mootha, V. K., Mukherjee, S., Ebert, B. L., Gillette, M. A., Paulovich, A., Pomeroy, S. L., Golub, T. R., Lander, E. S., et al. (2005) Gene set enrichment analysis: a knowledge-based approach for interpreting genome-wide expression profiles. Proceedings of the National Academy of Sciences, 102(43), 15545–15550.

8. Smyth, G. K. (2005) Limma: linear models for microarray data. In Bioinformatics and computational biology solutions using R and Bioconductor pp. 397–420 Springer.

9. Ritchie, M. E., Phipson, B., Wu, D., Hu, Y., Law, C. W., Shi, W., and Smyth, G. K. (2015) limma powers differential expression analyses for RNA-sequencing and microarray studies. Nucleic acids research, 43(7), e47–e47.

10. Love, M. I., Huber, W., and Anders, S. (2014) Moderated estimation of fold change and dispersion for RNA-seq data with DESeq2. Genome biology, 15(12), 1–21.

11. Satija, R., Farrell, J. A., Gennert, D., Schier, A. F., and Regev, A. (2015) Spatial reconstruction of single-cell gene expression data. Nature biotechnology, 33(5), 495–502.

12. Zeleznik, R., Foldyna, B., Eslami, P., Weiss, J., Alexander, I., Taron, J., Parmar, C., Alvi, R. M., Banerji, D., Uno, M., et al. (2021) Deep convolutional neural networks to predict cardiovascular risk from computed tomography. Nature communications, 12(1), 1–9.

13. Yao, D., Zhi-li, Z., Xiao-feng, Z., Wei, C., Fang, H., Yao-ming, C., and Cai, W.-W. (2022) Deep hybrid: multi-graph neural network collaboration for hyperspectral image classification. Defence Technology,.

14. Gaur, L., Bhatia, U., Jhanjhi, N., Muhammad, G., and Masud, M. (2021) Medical image-based detection of COVID-19 using deep convolution neural networks. Multimedia systems, pp. 1–10.

15. Miles, C., Bohrdt, A., Wu, R., Chiu, C., Xu, M., Ji, G., Greiner, M., Weinberger, K. Q., Demler, E., and Kim, E.-A. (2021) Correlator convolutional neural networks as an interpretable architecture for image-like quantum matter data. Nature Communications, 12(1), 1–7.

16. Sharma, P. K., Bisht, I., and Sur, A. (2021) Wavelength-based attributed deep neural network for underwater image restoration. ACM Journal of the ACM (JACM),.

17. Aliper, A., Plis, S., Artemov, A., Ulloa, A., Mamoshina, P., and Zhavoronkov, A. (2016) Deep learning applications for predicting pharmacological properties of drugs and drug repurposing using transcriptomic data. Molecular pharmaceutics, 13(7), 2524–2530.

18. Luo, Q., Mo, S., Xue, Y., Zhang, X., Gu, Y., Wu, L., Zhang, J., Sun, L., Liu, M., and Hu, Y. (2021) Novel deep learning-based transcriptome data analysis for drug-drug interaction prediction with an application in diabetes. BMC bioinformatics, 22(1), 1–15.

19. Hong, J., Hachem, L. D., and Fehlings, M. G. (2022) A deep learning model to classify neoplastic state and tissue origin from transcriptomic data. Scientific reports, 12(1), 1–7.

20. Chen, H.-I. H., Chiu, Y.-C., Zhang, T., Zhang, S., Huang, Y., and Chen, Y. (2018) GSAE: an autoencoder with embedded gene-set nodes for genomics functional characterization. BMC systems biology, 12(8), 45–57.

21. Liou, C.-Y., Cheng, W.-C., Liou, J.-W., and Liou, D.-R. (2014) Autoencoder for words. Neurocomputing, 139, 84–96.

22. Way, G. P. and Greene, C. S. (2018) Extracting a biologically relevant latent space from cancer transcriptomes with variational autoencoders. In PACIFIC SYMPOSIUM ON BIOCOMPUTING 2018: Proceedings of the Pacific Symposium World Scientific pp. 80–91.

23. Lopez, R., Regier, J., Cole, M. B., Jordan, M. I., and Yosef, N. (December, 2018) Deep generative modeling for single-cell transcriptomics. Nature Methods, 15(12), 1053–1058 Number: 12 Publisher: Nature Publishing Group.

24. Ding, J., Condon, A., and Shah, S. P. (May, 2018) Interpretable dimensionality reduction of single cell transcriptome data with deep generative models. Nature Communications, 9(1), 2002 Number: 1 Publisher: Nature Publishing Group.

25. Zhao, Y., Cai, H., Zhang, Z., Tang, J., and Li, Y. (September, 2021) Learning interpretable cellular and gene signature embeddings from single-cell transcriptomic data. Nature Communications, 12(1), 5261 Number: 1 Publisher: Nature Publishing Group.

26. Lotfollahi, M., Rybakov, S., Hrovatin, K., Hediyeh-zadeh, S., Talavera-López, C., Misharin, A. V., and Theis, F. J. (February, 2023) Biologically informed deep learning to query gene programs in single-cell atlases. Nature Cell Biology, 25(2), 337–350 Number: 2 Publisher: Nature Publishing Group.

27. Doersch, C. (2016) Tutorial on variational autoencoders. arXiv preprint arXiv:1606.05908,.

28. Higgins, I., Matthey, L., Pal, A., Burgess, C., Glorot, X., Botvinick, M., Mohamed, S., and Lerchner, A. (2016) beta-vae: Learning basic visual concepts with a constrained variational framework.

29. Rumelhart, D. E., Hinton, G. E., and Williams, R. J., Learning internal representations by error propagation. Technical report, California Univ San Diego La Jolla Inst for Cognitive Science (1985).

30. Torrente, A. A comprehensive human expression map.

31. McCall, M. N., Bolstad, B. M., and Irizarry, R. A. (2010) Frozen robust multiarray analysis (fRMA). Biostatistics, 11(2), 242–253.

32. McCall, M. N., Uppal, K., Jaffee, H. A., Zilliox, M. J., and Irizarry, R. A. (2011) The Gene Expression Barcode: leveraging public data repositories to begin cataloging the human and murine transcriptomes. Nucleic acids research, 39(uppl 1), D1011–D1015.

33. Margus, L., Wolfgang, H., et al. (2011) Assessing affymetrix GeneChip microarray quality. BMC.

34. McCall, M. N., Jaffee, H. A., and Irizarry, R. A. (2012) fRMA ST: frozen robust multiarray analysis for Affymetrix Exon and Gene ST arrays. Bioinformatics, 28(23), 3153–3154.

35. Liberzon, A., Birger, C., Thorvaldsdóttir, H., Ghandi, M., Mesirov, J. P., and Tamayo, P. (2015) The molecular signatures database hallmark gene set collection. Cell systems, 1(6), 417–425.

36. Pedregosa, F., Varoquaux, G., Gramfort, A., Michel, V., Thirion, B., Grisel, O., Blondel, M., Prettenhofer, P., Weiss, R., Dubourg, V., Vanderplas, J., Passos, A., Cournapeau, D., Brucher, M., Perrot, M., and Duchesnay, E. (2011) Scikit-learn: Machine Learning in Python. Journal of Machine Learning Research, 12, 2825–2830.

37. Traag, V. A., Waltman, L., and Van Eck, N. J. (2019) From Louvain to Leiden: guaranteeing well-connected communities. Scientific reports, 9(1), 5233.

38. Asperti, A. and Trentin, M. (2020) Balancing reconstruction error and Kullback-Leibler divergence in Variational Autoencoders. IEEE Access, 8, 199440–199448.

39. Qi, Y. and Fu, J. (2017) Research on the coagulation function changes in non small cell lung cancer patients and analysis of their correlation with metastasis and survival. J buon, 22(2), 462–467.

40. Sotiropoulos, G. P., Dalamaga, M., Antonakos, G., Marinou, I., Vogiatzakis, E., Kotopouli, M., Karampela, I., Christodoulatos, G. S., Lekka, A., and Papavassiliou, A. G. (2018) Chemerin as a biomarker at the intersection of inflammation, chemotaxis, coagulation, fibrinolysis and metabolism in resectable non-small cell lung cancer. Lung Cancer, 125, 291–299.

41. Gabazza, E. C., Taguchi, O., Yamakami, T., Machishi, M., Ibata, H., and Suzuki, S. (1993) Correlation between increased granulocyte elastase release and activation of blood coagulation in patients with lung cancer. Cancer, 72(7), 2134–2140.

42. Gezelius, E., Flou Kristensen, A., Bendahl, P., Hisada, Y., Risom Kristensen, S., Ek, L., Bergman, B., Wallberg, M., Falkmer, U., Mackman, N., et al. (2018) Coagulation biomarkers and prediction of venous thromboembolism and survival in small cell lung cancer: A sub-study of RASTEN-A randomized trial with low molecular weight heparin. PLoS One, 13(11), e0207387.

